# CAMITAX: Taxon labels for microbial genomes

**DOI:** 10.1101/532473

**Authors:** Andreas Bremges, Adrian Fritz, Alice C. Mchardy

## Abstract

The number of microbial genome sequences is growing exponentially, also thanks to recent advances in recovering complete or near-complete genomes from metagenomes and single cells. Assigning reliable taxon labels to genomes is key and often a prerequisite for downstream analyses. We introduce CAMITAX, a scalable and reproducible workflow for the taxonomic labelling of microbial genomes recovered from isolates, single cells, and metagenomes. CAMITAX combines genome distance-, 16S rRNA gene-, and gene homology-based taxonomic assignments with phylogenetic placement. It uses Nextflow to orchestrate reference databases and software containers, and thus combines ease of installation and use with computational reproducibility. We evaluated the method on several hundred metagenome-assembled genomes with high-quality taxonomic annotations from the TARA Oceans project, and show that the ensemble classification method in CAMITAX improved on all individual methods across tested ranks. While we initially developed CAMITAX to aid the Critical Assessment of Metagenome Interpretation (CAMI) initiative, it evolved into a comprehensive software to reliably assign taxon labels to microbial genomes. CAMITAX is available under the Apache License 2.0 at: https://github.com/CAMI-challenge/CAMITAX

## INTRODUCTION

The direct costs for sequencing a microbial genome are at an all-time low: a high-quality draft now costs less than $100, a “finished” genome sequence less than $500. This resulted in many culture-dependent genome studies, in which thousands of isolates—selected by e.g. their distinct phylogeny [1, 2], abundance in the human microbiome [3, 4], or biotechnological relevance [5, 6]—are sequenced.

Single cell genome and shotgun metagenome studies further contribute to this expansion in genome numbers by enabling access to the genome sequences of (as-yet) uncultured microbes [7–9]. Notably, new bioinformatics methods can reconstruct complete or near-complete genomes even from complex environments [10, 11], and easily scale to hundreds or even thousands of metagenome samples [12–16].

Typically, the sequencing and assembly of a new genome is merely a prerequisite for further bioinformatics analyses (and their experimental validation) to uncover novel biological insights by e.g. functional annotation [17, 18] or phenotype prediction [19, 20], which often require the genome’s taxonomy.

Historically, a bacterial or archaeal species was defined as a collection of strains that share one (or more) trait(s) and show DNA-DNA reassociation values of 70% or higher [21]. However, with the advent of genomics and—more recently—culture-independent methods, this definition was found to be impractical and difficult to implement [22].

Today, 16S rRNA gene similarity, average nucleotide identity (ANI), genome phylogeny, or gene-centric voting schemes are used for taxonomic assignments [23–28]. These approaches all have their merits (see below), but, to the best of our knowledge, no unifying workflow implementation existed. To jointly use these complementary approaches, we developed CAMITAX, a scalable and reproducible workflow that combines genome distance-, 16S rRNA gene-, and gene homology-based taxonomic assignments with phylogenetic placement onto a fixed reference tree to reliably infer genome taxonomy.

## METHODS

In the following, we describe CAMITAX’s assignment strategies and its implementation (Figure 1).

**Fig. 1.**
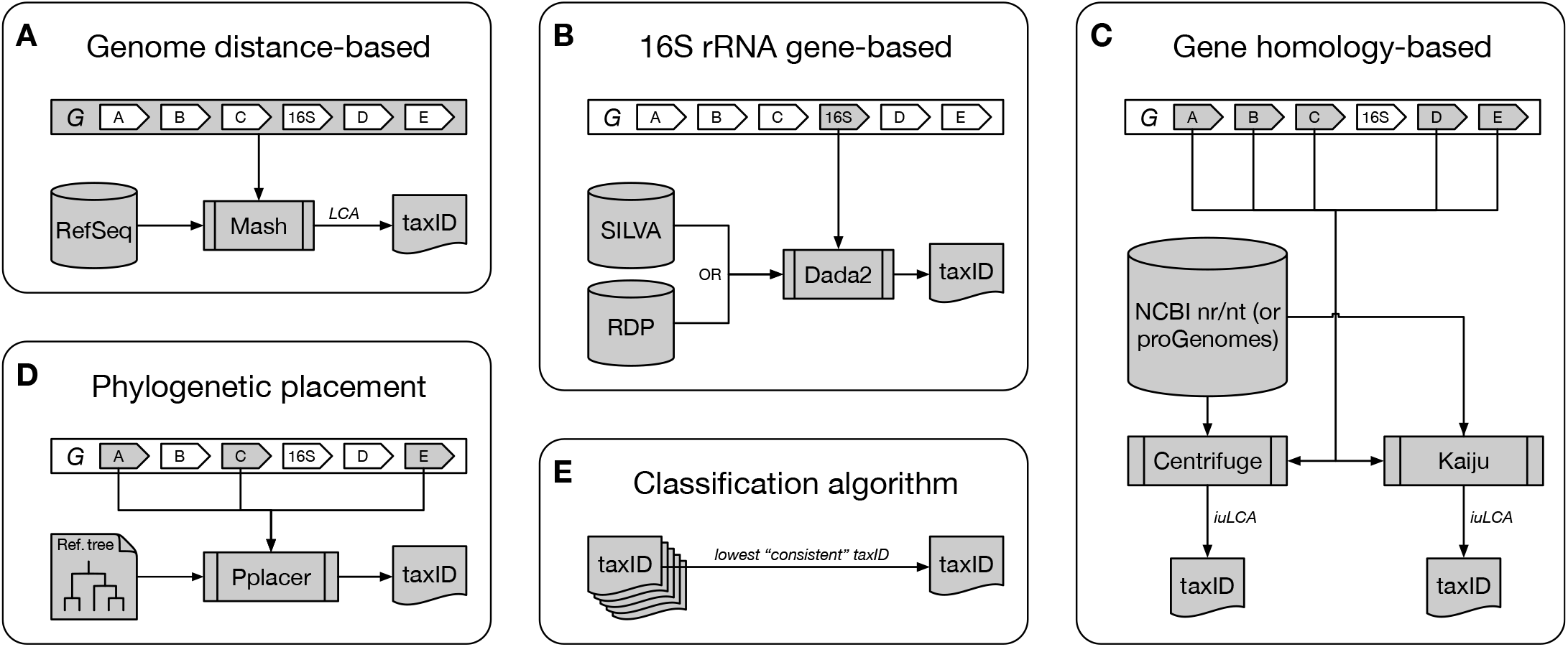
The CAMITAX taxonomic assignment workflow. CAMITAX assigns one NCBI Taxonomy ID (taxID) to an input genome *G* by combining genome distance-, 16S rRNA gene-, and gene homology-based taxonomic assignments with phylogenetic placement. **(A) Genome distance-based assignment.** CAMITAX uses Mash to estimate the average nucleotide identity (ANI) between *G* and more than a hundred thousand microbial genomes in RefSeq, and assigns the lowest common ancestor (LCA) of genomes showing >95% ANI, which was found to be a clear species boundary. **(B) 16S rRNA gene-based assignment.** CAMITAX uses Dada2 to label *G*’s 16S rRNA gene sequences using the naïve Bayesian classifier method to assign taxonomy across multiple ranks (down to genus level), and exact sequence matching for species-level assignments, against the SILVA or RDP database. **(C) Gene homology-based assignments.** CAMITAX uses Centrifuge and Kaiju to perform gene homology searches against nucleotide and amino acid sequences in NCBI’s nr and nt (or proGenomes’ genes and proteins datasets), respectively. CAMITAX determines the interval-union LCA (iuLCA) of gene-level assignments and places *G* on the lowest taxonomic node with at least 50% coverage **(D) Phylogenetic placement.** CAMITAX uses Pplacer to place *G* onto a fixed reference tree, as implemented in CheckM, and estimates genome completeness and contamination using lineage-specific marker genes. **(E) Classification algorithm.** CAMITAX considers the lowest consistent assignment as the longest unambiguous root-to-node path in the taxonomic tree spanned by the five taxIDs derived in (A)–(D), i.e. it retains the most specific, yet consistent taxonomic label among all tools.

### Genome distance-based assignment

An ANI value of 95% roughly corresponds to a 70% DNA-DNA reassociation value (the historical species definition) [24]. In other words, strains from the same species are expected to show >95% ANI [29]. This species boundary appears to be widely applicable and has been confirmed in a recent large-scale study, in which the analyses of 8 billion genome pairs revealed a clear genetic discontinuity among known genomes, with 99.8% of the pairs showing either >95% intra-species ANI or <83% inter-species ANI values [30].

CAMITAX uses Mash [31] to rapidly estimate the input genomes’ ANI to all bacterial or archaeal genomes in the RefSeq database [32] (114,176 strains as of 2018-05-10). CAMITAX’s genome distance-based assignment is the lowest common ancestor (LCA) of all Mash hits with >95% ANI; a genome is placed at *root* if there is no RefSeq genome with >95% ANI.

This strategy works best if the query genome is more than 80% complete (Mash does not accurately estimate the genome-wide ANI of incomplete genomes [33]) and is represented in RefSeq. CAMITAX’s other assignment strategies are complementary by design and better suited for incomplete genomes or underrepresented lineages. If a Mash hit is found, however, CAMITAX most likely assigns a taxonomy at the species or genus level.

### 16S rRNA gene-based assignment

The 16S rRNA gene is widely used for classification tasks because it is an universal marker gene likely present in all bacteria and archaea [34, 35].

CAMITAX uses nhmmer [36] to identify 16S rRNA genes in the input genomes and Dada2 [37] to assign taxonomy. Dada2 uses the naïve Bayesian classifier method [38] for kingdom to genus assignments, and exact sequence matching against a reference database for species assignments. CAMITAX supports two commonly used databases: SILVA [39] and RDP [40], which both were found to map back well to the NCBI Taxonomy [41].

Of course, this strategy only is applicable if the genome assembly contains a copy of the 16S rRNA gene—which is not always the case, particularly for genomes recovered from metagenomes or single cells.

### Gene homology-based assignments

Metagenomics and single cell genomics are complementary approaches providing access to the genomes of (as-yet) uncultured microbes, but both have strings attached: Single amplified genomes (SAGs) suffer from amplification bias and, as a consequence, are often incomplete [42, 43]. Metagenome-assembled genomes (MAGs) on the other hand rarely contain full-length 16S rRNA genes [44, 45]. While there are notable exceptions to this rule [46, 47], the above assignment strategies are generally not expected to work well for today’s SAGs and MAGs.

To overcome these problems, CAMITAX implements a gene-based voting scheme. It uses Prodigal [48] to predict protein-coding genes, and then Centrifuge [49] and Kaiju [50] for gene homology searches on the nucleotide and protein level, respectively. Both tools scale to large reference databases, such as NCBI’s nr/nt [51], but (by default) CAMITAX resorts to the (much smaller) proGenomes genes and proteins datasets [52, 53]. The proGenomes database was designed as a resource for consistent taxonomic annotations of bacteria and archaea.

Inferring genome taxonomy from a set of gene-level assignments is not trivial, and—inspired by procedures implemented in anvi’o [27] and dRep [33]—CAMITAX places the query genome on the lowest taxonomic node with at least 50% support in gene assignments (which corresponds to the interval-union LCA algorithm [28]) for nucleotide and protein searches.

### Phylogenetic placement

CAMITAX uses CheckM [25] for a phylogeny-driven estimate of taxonomy. Relying on 43 phylogenetically informative marker genes (consisting primarily of ribosomal proteins and RNA polymerase domains), CheckM places the query genome onto a fixed reference tree with Pplacer [54] to infer taxonomy. We note that phylogenetic placement is often quite conservative and does not necessarily provide resolution at the species level [26, 55].

Lastly, CAMITAX reports the query genome’s completeness and contamination as estimated by CheckM using its lineage-specific marker genes [25].

### Classification algorithm

CAMITAX considers the lowest consistent assignment as the longest unambiguous root-to-node path in the taxonomic tree spanned by the individual assignments, i.e. it retains the most specific, yet consistent taxonomic label among all tools. For example, CAMITAX would determine as “consistent” assignments for the individual assignments (derived with the different assignment strategies) the following:

- 3× *E. coli*, 2× *Bacteria* ↦ *E. coli*
- 3× *E. coli*, 2× *E. albertii* ↦ *Escherichia*
- 3× *E. coli*, 2× *Archaea* ↦ Root

This strategy is more robust than computing the lowest common ancestor (LCA) of individual assignments because outliers, e.g. missing predictions of conservative methods, don’t affect the overall assignment.

At the same time, requiring a consistent assignment is less error-prone than e.g. selecting the maximal root-to-leaf path, which would introduce many false-positive assignments especially on lower ranks.

### Implementation

CAMITAX incorporates many state-of-the-art pieces of software, and automatically resolves all software and database dependencies with Nextflow [56] in a containerized environment (Table 1). This fosters reproducibility in bioinformatics research [57, 58], and we strongly suggest to run CAMITAX using BioContainers [59] (automated container builds for software in bioconda [60]). CAMITAX can be run on a local machine or in a distributed fashion.

**Table 1.** Software used in the CAMITAX workflow.

| Software | Version | BioContainer |
| --- | --- | --- |
| Centrifuge | 1.0.3 | centrifuge:1.0.3-py36pl5.22.0_2 |
| CheckM | 1.0.11 | checkm-genome:1.0.11-0 |
| Dada2 | 1.6.0 | bioconductor-dada2:1.6.0-r3.4.1_0 |
| Kaiju | 1.6.2 | kaiju:1.6.2-pl5.22.0_0 |
| Mash | 2.0 | mash:2.0-gsl2.2_2 |
| Nhmmer | 3.1 | - |
| Pplacer | 1.1 | - |
| Prodigal | 2.6.3 | prodigal:2.6.3-0 |

CAMITAX automatically resolves all software dependencies with Nextflow using BioContainers in a containerized environment. Nhmmer and Pplacer are bundled with CheckM.

## RESULTS

We applied CAMITAX to real data not present in its databases, a recent collection of 885 bacterial and archaeal MAGs from Delmont *et al.* [15], who used state-of-the-art metagenomic assembly, binning, and curation strategies to create a non-redundant database of microbial population genomes from the TARA Oceans project [61].

Delmont *et al.* used CheckM for an initial taxonomic inference of the MAGs. Thereafter, they used Centrifuge [49], RAST [62], and manual BLAST searches of single-copy core genes against NCBI’s nr/nt to manually refine their taxonomic inferences. Lastly, they trained a novel machine learning classifier to also identify MAGs affiliated to the Candidate Phyla Radiation (CPR) [8].

As expected, CAMITAX outperformed CheckM, which is rather conservative in its assignments, by adding low-ranking annotations based on high-quality predictions of other tools, such as Kaiju (Figure 2). Notably, 95% of CAMITAX’s predictions were consistent with Delmont *et al.*, i.e. the two assignments were on the same taxonomic lineage and their LCA is either of the two. CAMITAX assignments of 46 MAGs (5%) were in conflict with the manually curated taxonomy. Of these, CAMITAX made species assignments for twelve MAGs based on Mash hits against RefSeq genomes. These we consider trustworthy because >95% ANI was shown to be a clear species boundary [30], and we assume that Delmont *et al.* assigned them incorrectly. On the other hand, CAMITAX for instance misclassified MAGs affiliated to the Candidate Phyla Radiation based on their 16S rRNA gene sequences to other phyla.

**Fig. 2.**
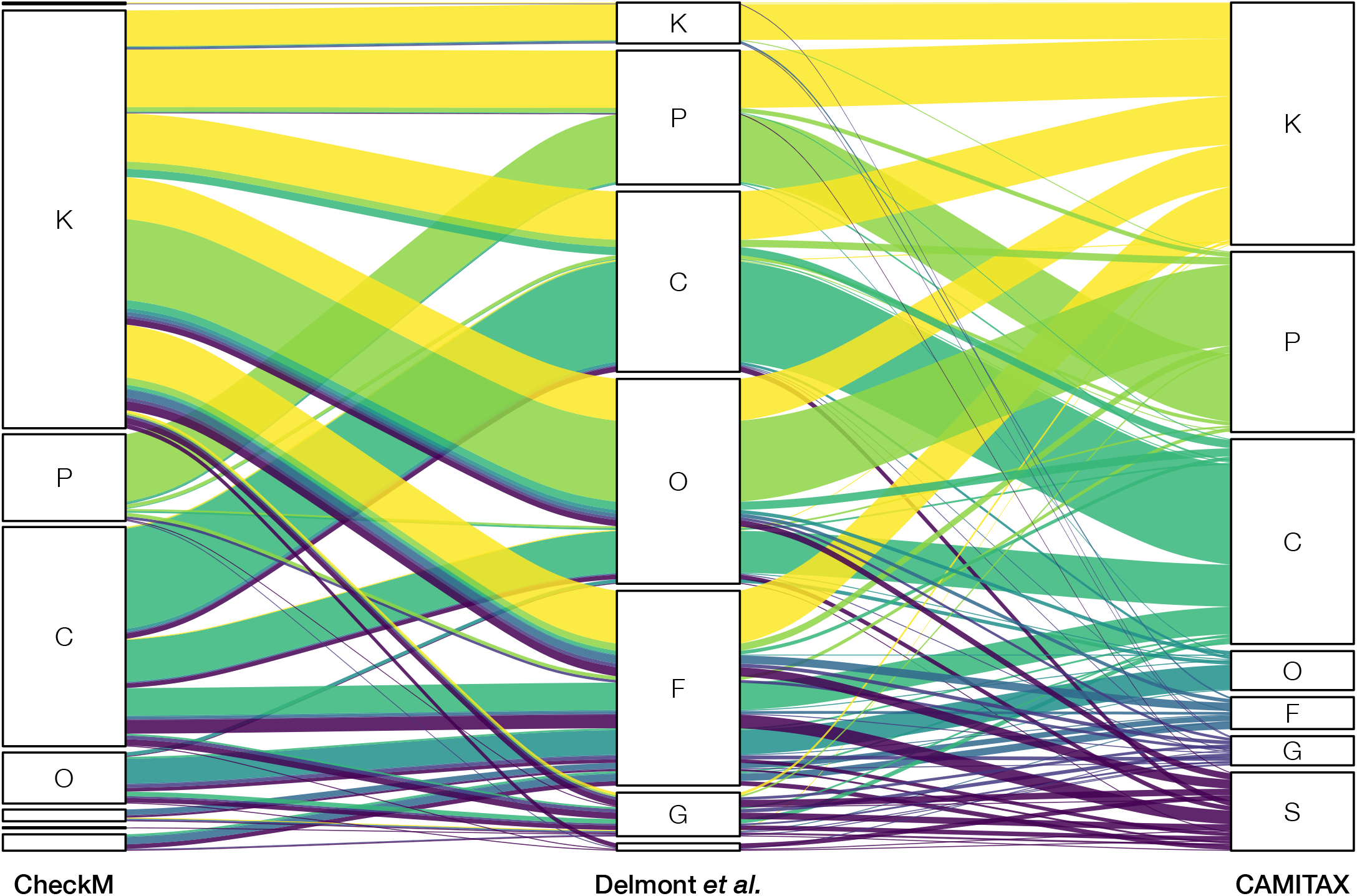
Comparison of high quality taxonomic assignments for 885 MAGs. Using genome-resolved metagenomics, Delmont *et al.* assembled 885 bacterial and archaeal genomes from the TARA Oceans metagenomes and used CheckM for an initial taxonomic inference. Subsequently, they manually refined the taxonomic assignments using additional analyses and expert knowledge. The alluvial diagram shows the assigned taxonomic ranks for CheckM (left), manual curation (middle), and CAMITAX (right) on kingdom, phylum, class, order, family, genus, and species level.

To quantify taxonomic assignment performance, we calculated precision, recall, and accuracy across all ranks with AMBER 2.0 [63] (Figure 3). As the gold standard, we used the Delmont *et al.* assignments up to genus rank. CAMITAX was very precise down to class level and reasonably (>80%) precise below. Overall, it was more accurate across all ranks than each of its assignment strategies individually. While the recall of CAMITAX dropped at the mid-range ranks, it recovered for genus level assignments.

**Fig. 3.**
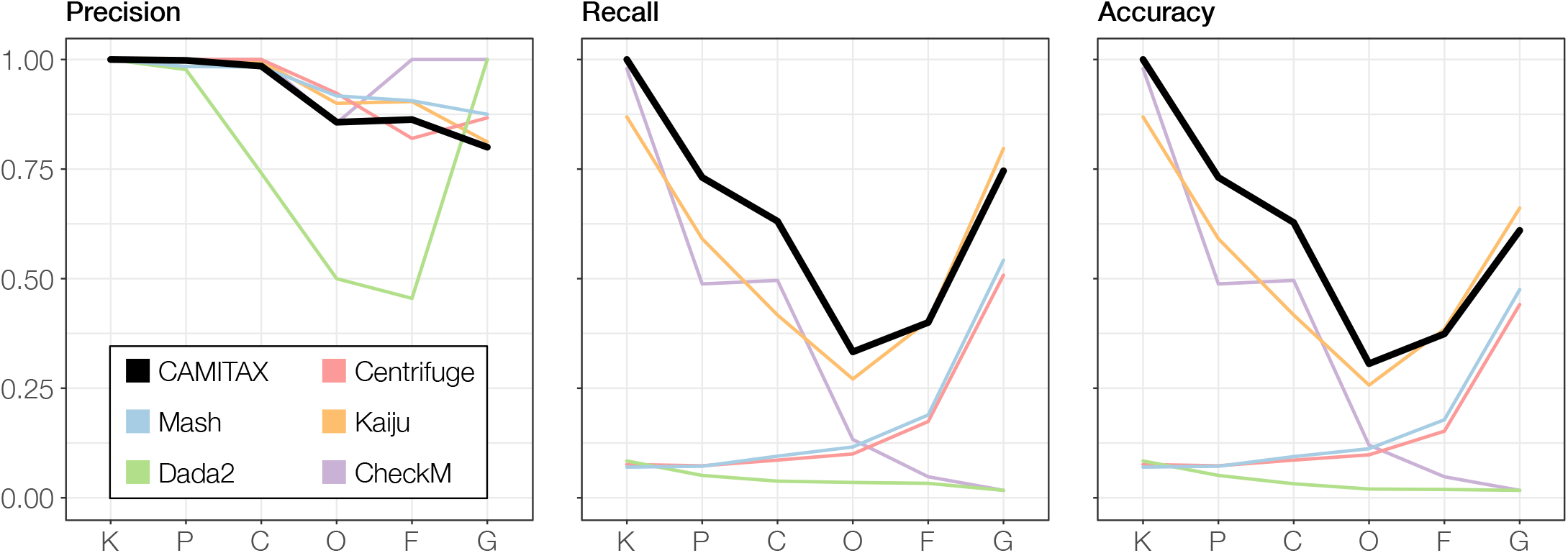
Taxonomic assignment performance metrics across ranks for 885 MAGs. The manually assigned taxonomy by Delmont *et al.* is used as the gold standard to benchmark against. Shown are precision, recall, and accuracy for CAMITAX (and the individual tools combined therein) on kingdom, phylum, class, order, family, and genus level.

We thus propose CAMITAX as a reliable and reproducible taxonomic assignment workflow, ideally followed by a manual refinement step—as always.

## DISCUSSION

CAMITAX was initially developed while preparing the second Critical Assessment of Metagenome Interpretation (CAMI) challenge [64]. The challenge datasets include new genomes from taxa (at different evolutionary distances) not found in public databases yet, which need high quality taxon labels for the subsequent microbial community and metagenome data simulation [65]. Due to this need, we created CAMITAX to systematically double-check, newly infer, or refine genome taxon label assignments in a fully reproducible way.

CAMITAX combines different taxonomic assignment strategies in one unifying workflow implementation. It uses Nextflow to orchestrate reference databases and software containers. Therefore, both databases and software can be easily substituted, providing the flexibility to cope with rapid change of standards oftentimes observed in the field. For instance, Parks *et al.* recently proposed a standardized bacterial taxonomy based on genome phylogeny, the so-called Genome Taxonomy Database (GTDB) [66]. While CAMITAX currently uses the NCBI Taxonomy [67], it is (at least in principle) agnostic to the underlying database and could thus be easily adapted to other taxonomy versions that will arise in future.

## SOFTWARE AND DATA AVAILABILITY

CAMITAX is implemented in Nextflow and Python 3, and is freely available under the Apache License 2.0 at https://github.com/CAMI-challenge/CAMITAX.

Mash sketches for all bacterial and archaeal genomes in RefSeq, snapshots of the NCBI Taxonomy databases, and Centrifuge and Kaiju indices for the proGenomes genes and proteins datasets, respectively, are collected under doi:10.5281/zenodo.1250043. The snapshots used in this study, generated on 2018-05-10, are available under doi:10.5281/zenodo.1250044.

Dada2-formatted training fasta files, derived from SILVA (release 132) and RDP (training set 16, release 11.5), are available under doi:10.5281/zenodo.1172782 and doi:10.5281/zenodo.801827, respectively.

Lastly, the CheckM reference databases are available at https://data.ace.uq.edu.au/public/CheckM_databases.

## AUTHORS’ CONTRIBUTIONS

AB implemented the software, performed experiments, and wrote the paper with comments from AF and ACM. AF thoroughly tested the software. AB and ACM jointly conceived the project and evaluated results. All authors read and approved the final manuscript.

## ACKNOWLEDGEMENTS

The authors thank Peter Belmann for Nextflow and Docker tips, Fernando Meyer for early beta testing, and the Isaac Newton Institute for Mathematical Sciences for its hospitality during the programme MTG, which was supported by EPSRC Grant Number EP/K032208/1.

